# Exophers are components of mammalian cell neurobiology in health and disease

**DOI:** 10.1101/2021.12.06.471479

**Authors:** Ibrar Siddique, Jing Di, Christopher K. Williams, Daniela Markovic, Harry V. Vinters, Gal Bitan

## Abstract

Maintenance of cellular homeostasis is critically important for the survival of cells and organisms. Degradation and recycling of biomolecules and whole organelles is an essential mechanism for maintaining cellular homeostasis. The main systems responsible for these processes are the ubiquitin-proteasome system and autophagy-lysosome pathway. Another mechanism was reported in *c. elegans*—formation of large, membrane-enclosed projections called “exophers,” into which cells direct debris and toxic protein aggregates^1^. Exophers were shown to act as large, temporary disposal compartments that disconnected from the cells within several hours. Here, we report the discovery of exophers in the mammalian brain, including the brains of humans and mice. Similar to those described in nematodes, the mammalian exophers appear to emanate from the cell body, initially connected by a nanotube, and eventually disconnect. Rare, innate exophers were found in healthy human brain and primary neurons from wild-type mice, presumably mediating transfer of large cargo between cells. The number of exophers increased as an adaptive response under proteostatic stress, e.g., in Alzheimer’s disease brain or in primary neurons from two tauopathy mouse models, where the exophers likely facilitated expulsion of proteotoxic material. Our findings suggest that exopherogenesis is a rare, innate house-keeping process that is elevated adaptively in response to proteostatic pressure and is a conserved mechanism from nematodes to humans.

## Introduction

Cells are dynamic environments where biomolecules constantly are recycled and replaced by newly synthesized copies. This turnover process ensures a constant supply of new, functional nucleic acids, proteins, lipids, and sugars, and degradation of old and/or damaged biomolecules and organelles. Eukaryotic cells rely primarily on the ubiquitin-proteasome system (UPS) and the autophagy-lysosome pathway (ALP) for protein degradation. In the UPS, proteins are marked for proteasomal degradation by conjugation of ubiquitin^2^, whereas in the ALP, molecules and organelles destined for recycling are encapsulated first by autophagosomes, which fuse with lysosomes in which enzymatic degradation occurs^3^. The UPS degrades primarily short-lived proteins and soluble misfolded proteins, and has an important role in cell signaling, transcription, cell proliferation and other vital cellular pathways^4^. The ALP’s main role is clearance of long-lived proteins, insoluble protein aggregates, whole organelles (e.g., mitochondria), and intracellular invaders^5^, and it participates in crucial adaptive responses to various proteostasis stresses, such as nutrient deprivation, hypoxia, and oxidative stress^6^.

In 2017, a novel additional mechanism allowing cells to handle proteostatic stress was described by Melentijevic et al. who found that adult *c. elegans* neurons extrude large (~4 μm) membrane-enclosed vesicle-like structures they named exophers^1^. Exophers were shown to emanate from neurons, initially connected by a nanotube and eventually disconnect, leaving the exopher separate from the originating neuron. The exophers were demonstrated to contain protein aggregates and organelles and their production was enhanced in the nematodes by inhibition of chaperone expression, the UPS, or the ALP. Moreover, neurons that produced exophers showed increased survival compared to neurons that did not under proteotoxic conditions^1^. Follow-up studies in *c. elegans* showed that neuronal exopherogenesis also can be induced by nutrient deprivation and may be controlled by specific, non-cell autonomous signaling pathways related to lipid synthesis and/or the fibroblast growth factor/RAS/MAPK pathway^7^. Body-wall muscle cell exophers were observed in *c. elegans* to support embryo development^8^, and exophers of mouse and human cardiomyocytes also were reported to expel damaged mitochondria and other cargo to be taken up by macrophages^9^. However, it has been unknown if exophers exist in the mammalian brain, if they indeed assist in removal of cell debris under proteostatic stress as observed in *c. elegans*, and whether they also have a role in normal physiology, other than the recently reported nematode embryo support.

During the analysis of tauopathy mouse brains^10^, we noticed rare exophers, similar to those described in *c. elegans*, which were attached to neuronal bodies via nanotubes (Fig. 1). This discovery of exophers in the brain of a mammal prompted us to ask if similar structures could be found in the human brain and whether evidence supporting the role previously hypothesized for exophers as a supportive mechanism to the UPS and ALP^1,7^ under proteostatic stress could be found in humans or mice.

**Figure 1:**
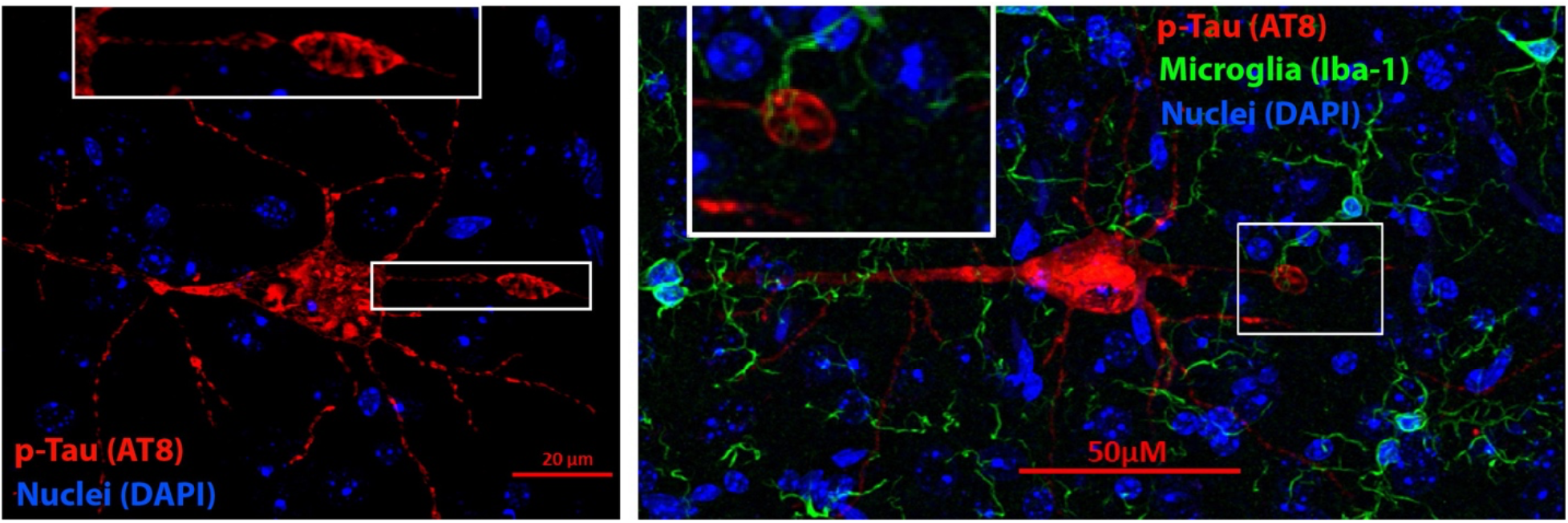
Exophers found in the brain of P301S-tau mice. Mouse hippocampal sections were stained with monoclonal antibody AT8 (red), which recognizes the phosphorylation sites at S202 and T205, the nuclear stain 4’,6-diamidino-2-phenylindole (DAPI, blue), and the microglia marker ionized calcium-binding adapter molecule 1 (Iba-1, green, right image only) and imaged using a confocal fluorescence microscope. Insets show zoomed in images of the exophers and connecting nanotubes. In the right image, the exopher appears to be in contact with microglia branches.

As a recently discovered cellular structure, there are no known markers for exophers. They have been defined morphologically as quasi-spherical bodies attached to, or detached from, the original neuron, smaller or of similar size to the cell body, and potentially containing multiple organelles. Exophers are relatively easy to spot when they are small compared to the cell body and the connecting nanotube is visible, but when they are approximately as large as the cell itself and have disconnected, it may be challenging to discern them from the cell bodies. Our initial observations in 8-months-old P301S-tau mice (Fig. 1), in agreement with the previous reports, showed that the most distinguishing feature is the absence of a nucleus in the exophers, which always stain negative with DAPI or Hoechst dyes. This distinction allowed us to look for exophers in human brains.

To visualize neuronal morphology, we stained hippocampi from two patients with Alzheimer’s disease, Braak stage VI, one patient with frontotemporal lobar degeneration (FTLD), one patient with progressive supranuclear palsy (PSP), and two elderly control (no neurologic disease) individuals (Table 1). We used a fluorescently labeled antibody against β-III tubulin, a ubiquitous neuronal marker^11^, allowing visualization of multiple exophers in these brains, including in the process of emanating from the cell body (Fig. 2a), connected via a nanotube (Fig. 2b), or disconnected (Fig. 2c). Exophers were detected in all the brains, though in the two control brains only a few exophers were found in the hippocampus, whereas > 20 exophers were found in similar total areas of each of the disease brains (Fig. 2d). Comparison of the density of exophers in the hippocampus showed a 2–3 times higher density in the disease brains than in the control brains (*p* = 0.0068) and was slightly higher in the FTLD and PSP than in the AD brains (Table 2). The exophers had a broad size distribution (size was measured as area in 2D images), ranging from 1.4 to 253.9 μm^2^. Most exophers were between 5 and 60 μm^2^ and the differences among the average sizes were statistically insignificant (Fig. 2d), yet exophers larger than 61 μm^2^ were found only in the disease brains.

**Figure 2:**
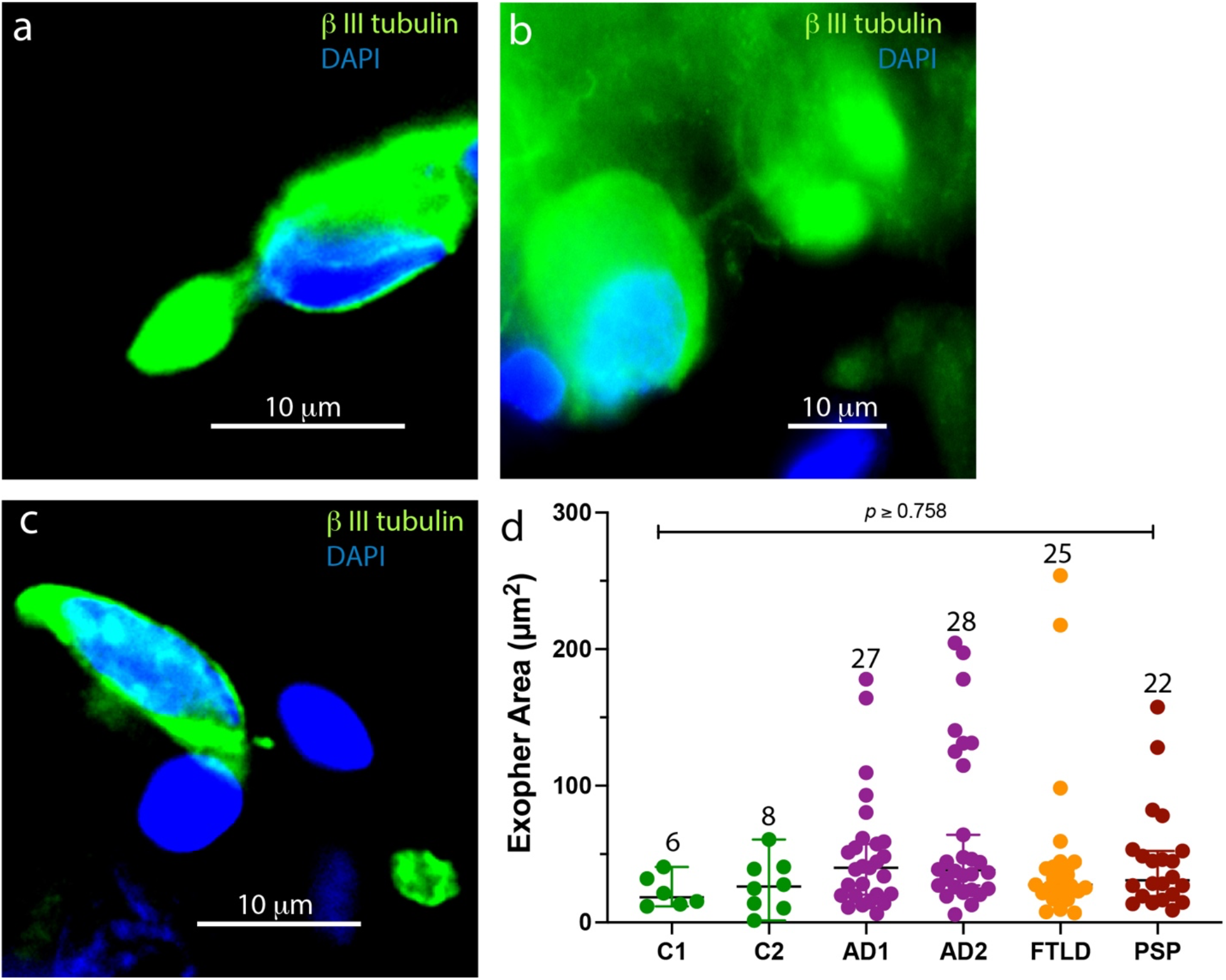
Human-brain exophers. Human hippocampal sections were stained with an anti-β-III tubulin antibody (green) to outline the morphology of the cells and exophers and DAPI (blue). a) An exopher connected directly to the cell body. b) An exopher connected via a nanotube. c) An exopher disconnected from the adjacent cell. d) Comparison of exopher areas in human brains. The data are presented as median ± CI. The number of exophers in each case is indicated above each column. *P*-values were calculated using a one-way ANOVA with post-hoc Tukey test.

**Table 1.** Human brains used in the study.

| **Table 1. Human brains used in the study** |  |  |
| --- | --- | --- |
| **Disease** | **Age** | **Sex** |
| Control 1 (cause of death: pneumonia) | 72 | M |
| Control 2 (cause of death: liver disease) | 69 | F |
| Alzheimer's Disease 1 (AD1), Braak stage VI | 75 | F |
| Alzheimer's Disease 2 (AD2), Braak stage VI | 99 | M |
| Progressive supranuclear palsy | 68 | M |
| Frontotemporal lobar degeneration | 79 | F |

**Table 2.** Exopher density in human hippocampus.

| **Table 2. Exopher density in human hippocampus** |  |  |  |  |
| --- | --- | --- | --- | --- |
| **Comparison** | **Rate Ratio** | **Lower Confidence Level** | **Upper Confidence Level** | **p-Value** |
| **AD vs Normal** | 2.12 | 1.12 | 4.02 | 0.0211 |
| **FTLD vs Normal** | 2.67 | 1.36 | 5.25 | 0.0136 |
| **PSP vs Normal** | 2.53 | 1.26 | 5.05 | 0.0175 |

The observation of exophers was not limited to using β-III tubulin. We observed similar structures in hippocampus sections stained with another neuronal marker, microtubule-associated protein 2 (MAP2, Supplemental Fig. 1a-c). The area of the 58 exophers observed using MAP-2 was similar to those in the β-III tubulin-stained sections (Supplemental Fig. 1d).

Interestingly, in some cases, small exophers were apparent midway between the cell body and main exophers, connected to both by nanotubes (Supplemental Fig. 2 and Supplemental Movie 1). These structures suggested that while the exophers still are connected to the parent cell, small “packages” of cellular material may be added to the main exophers, similar to a phenomenon described recently in *c. elegans* muscle-cell exophers^8^.

In AD, two offending proteins, Aβ and tau, aggregate in the brain, whereas in tau-related FTLD and PSP only tau aggregates are found in the brain, not aggregates of Aβ. As exophers were suggested to be a mechanism cells use to expel excess aggregated proteins, we hypothesized that exophers in the AD, FTLD, and PSP brains would contain hyperphosphorylated tau (p-tau), whereas only the AD brain exophers would contain also Aβ. To test the hypothesis, we stained the brain sections for β-III tubulin or MAP2, hyperphosphorylated tau, and Aβ42 (Supplemental Fig. 3). Indeed, we found p-tau staining in exophers of all four disease brain samples (Supplemental Fig. 3c-f), whereas only in the AD brain, Aβ42 was found in the exophers (Supplemental Fig. 3c, d). Neither Aβ nor p-tau were found in the control brains (Supplemental Fig. 3a, b). The number of p-tau- or Aβ42-positive exophers was higher than exophers staining negative for these proteins in all four disease brains (Supplemental. Fig. 4). Interestingly, the predicted least-square mean area of tau-positive exophers, 68.1 μm^2^ was larger (p = 0.0203, 2-way ANOVA) than that of tau-negative exophers, 34.3 μm^2^, whereas the predicted least-square mean area of Aβ42-positive (55.7 μm^2^) and Aβ-negative (50.1 μm^2^) exophers in the AD brain was similar (*p* = 0.7927, Supplemental. Fig. 4), raising the possibility that exophers may contribute to the prion-like cell-to-cell spreading of pathological forms of tau, but not Aβ42 in the brain.

To explore whether tauopathy-induced proteostatic stress causes temporal changes in exopher density and size, we used primary hippocampal neurons derived from two tauopathy mouse models: P301S-tau (PS19 line), expressing 1N4R MAPT containing an FTD-associated mutation leading to a P301S substitution, under control of the mouse PrP promoter^12^; and hTau, which expresses all 6 human tau isoforms under the human tau promoter on a mouse-tau-null background^13^. Neurons from wild-type mice were used as a control. Primary neurons gradually senesce after ~3 weeks in culture and therefore can serve as a proxy for cell-age-related and neurodegenerative changes^14,15^.

Cells were stained for MAP-2 and DAPI on day 7, 14, or 21 and analyzed together to maximize analysis consistency. We found exophers in the primary hippocampal cultures of all three mouse lines (Fig. 3a-c), yet the numbers were clearly higher in the tauopathy models than in the wildtype mice (Fig. 3d-f). The finding of exophers in wild-type neurons at the earliest time point, together with the rare exophers found in normal human brain, suggested that exophers are used as an innate housekeeping mechanism by neurons. Their increase with time and substantially higher numbers in the tauopathy models supported the previous observations in nematodes of upregulated neuronal exopherogenesis under proteostatic stress^1,7^.

**Figure 3:**
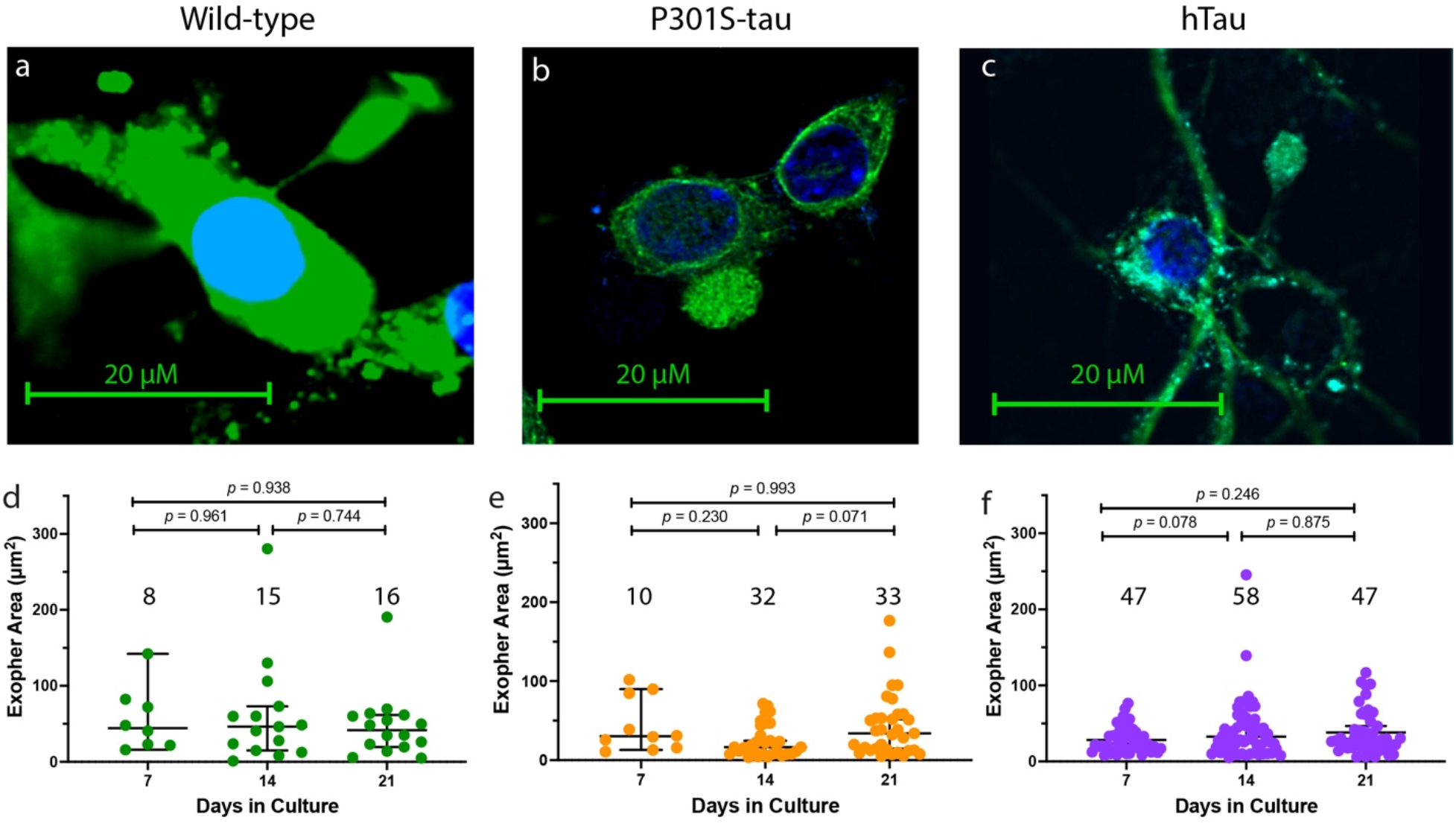
Exophers in wild-type, P301S-tau, and hTau mouse primary neurons. Primary hippocampal cultures of wild-type, P301S-tau, or hTau mice were analyzed on day 7, 14, or 21 in culture. a-c) Representative images of exophers in (a) wild-type mouse neurons (fluorescence microscope), (b) P301S-tau mouse neurons (confocal fluorescence microscope), and (c) hTau mouse neurons (confocal fluorescence microscope). d-f) Time-dependent exopher number and area in wild-type (d), P301S-tau (e), and hTau (f) mouse hippocampal neurons. The data are presented as median ± CI. The total number of exophers found at each time point is indicated above each column. *P*-values were calculated using a one-way ANOVA with post-hoc Tukey test.

The number of the exophers increased substantially between 7 and 14 days in the cultures of the wild-type (Fig. 3d) and P301S-tau mice (Fig. 3e) and changed little on day 21. The initial similar numbers in these two mouse lines suggest that proteostatic stress in the P301S-tau primary hippocampal neurons is minimal on day 7 in culture but increases substantially during the second week in culture, reaching approximately twice the number in the wild-type mice. In contrast, the number of exophers in the hTau hippocampal neurons was already high on day 7 (Fig. 3f) and changed minimally up to day 21, suggesting that proteostatic stress in this line starts earlier and is higher compared to the P301S-tau model.

The areas of the exophers varied widely, ranging from 2.1 to 280.3 μm^2^ (Fig. 3d-f). The median differences among the three mouse lines were small and changed minimally with cell age. The size range was similar to the exophers observed in the human brains (Fig. 2d) and clearly larger, on average, than the 11.3 μm^2^ average exopher area reported in *c. elegans*^1^, in agreement with the size difference between mammalian and nematode neurons.

To test whether the difference in exopher number in the primary hippocampal neurons between the two tauopathy mouse lines, and the increase in their number with time in culture correlated with tau accumulation, we stained the cells for MAP-2 to allow observation of exophers and for human tau using monoclonal antibody HT7 (Fig. 4). Again, we observed more exophers in the hTau hippocampal neurons compared to the P301S-tau neurons. However, this analysis revealed a unique behavior in each model. In the P301-tau neurons, about half of the relatively few exophers found on day 7 were tau positive (Fig. 4c). The number of tau-negative neurons changed little with culture age, whereas tau-positive exophers approximately tripled during the second week in culture and their number remained similar at 21 days, revealing that tau-positive exophers were responsible for the increase in exopher number shown in Fig. 3e. Overall, the predicted least-square mean area of tau-positive exophers (46.8 μm^2^) was larger than that of tau-negative exophers (25.6 μm^2^, *p* = 0.008, 2-way ANOVA).

**Figure 4:**
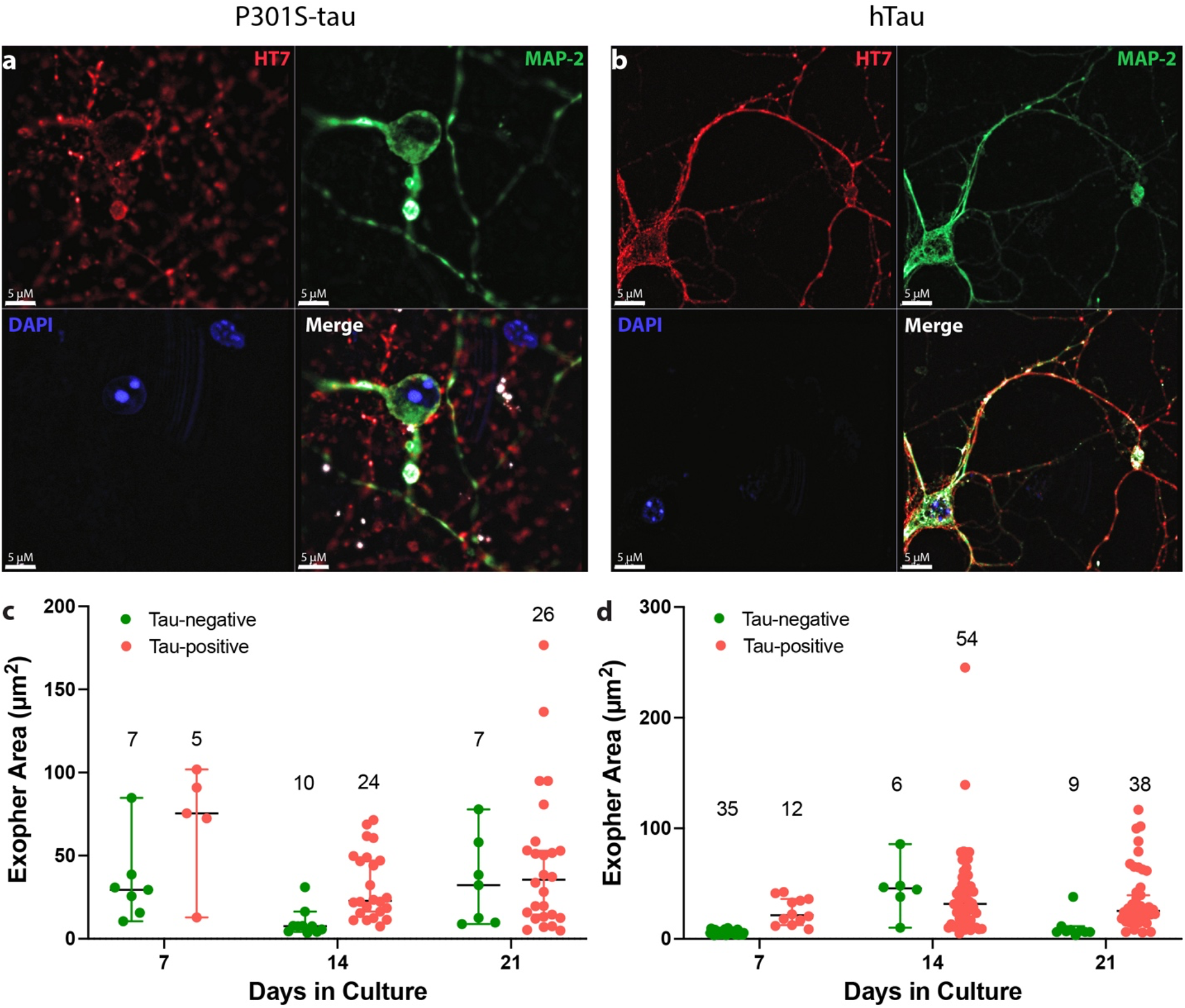
Tau-positive exophers tend to be larger than tau-negative exophers. Primary hippocampal cultures of P301S-tau (a) and hTau (b) mice were stained for MAP-2 (green) and DAPI (blue). The area of the exophers was measured using ImageJ and quantified separately for tau-negative and tau-positive exophers of P301S-tau (c) and hTau (d) mice on day 7, 14 and 21 in culture. The number of exophers observed at each time point is indicated above each column.

Interestingly, in the hTau neurons, on day 7, tau-positive exophers accounted for only a quarter of all exophers, whereas on days 14 and 21 the number of tau-positive exophers was 9-times, and ~4-times that of tau-negative exophers. Tau-negative exophers were mainly small on day 7 (median = 5.5 μm^2^) and day 21 (median = 6.9 μm^2^), and considerably larger on day 14 (median = 45.7 μm^2^). In comparison, there was relatively little variation in the median size of tau-positive exophers (21.3, 31.7, and 25.5 μm^2^ on days 7, 14, and 21, respectively). This distinct behavior suggests that the neurons may be stressed due to the presence of human tau early on, yet the reason for the different temporal dependence of exopher area on presence of tau is unclear. One week later, on day 14, both the number and the size of the exophers increase substantially and most of the exophers are tau-positive, whereas on day 21 the number and size decrease, possibly reflecting a decline in the culture’s health in response to tau’s proteotoxicity. Overall, the analysis of tau-negative and tau-positive exophers suggests that P301S-tau neurons have mostly a low level of innate exophers, increasing the number and size as an adaptive response on day 14 and remaining stable on day 21, whereas in the hTau neurons, an adaptive response is observed already on day 7, increases dramatically on day 14, and shows a decline as the culture ages on day 21.

Whereas the role of adaptive exophers, removing excess toxic protein aggregates from the neuron body in response to proteotoxicity, is relatively easy to understand, the physiologic role of the putative innate exophers is not known. Because cells transfer content into the exophers apparently using nanotubes, and nanotubes also are known to facilitate transfer of cargo between cells^16^, we hypothesized that innate exophers might be used to facilitate intercellular communication, particularly for transfer of cargos larger than would fit in a nanotube. Because exophers are rare and currently are defined mainly morphologically, to further explore their content and function, we sought a simpler system where they could be expected to be found reproducibly and chose the SH-SY5Y human neuroblastoma cell line as such a system.

Unlike in the human brain or primary cultures, here we sought, in particular, exophers connected to nanotubes, which are difficult to observe due to their narrow width. We found in these experiments that brightfield images at 60X magnification were best for visualizing the nanotube-connected exophers. We encountered approximately one exopher per chamber under normal conditions and analyzed 125 exophers across multiple experiments. The exophers were similar in morphology (Supplemental Fig. 5a, b) and size distribution (Supplemental Fig. 5c) to those we identified previously in human brain sections and mouse primary neurons. We initially analyzed 62 exophers in this system, whose area ranged from 1.5 to 204.1 μm^2^.

Close inspection of exophers in the SH-SY5Y cells yielded new observations: In a few cases, we observed an exopher emanating from another exopher (Figure 5a, b). In other cases, one exopher appeared to be connected via nanotubes to two or more cells (Figure 5c, d). In rare cases, two exophers originating in two different cells appeared to be in contact with each other (Figure 5e, f). These observations support the hypothesis that exophers may be used to facilitate cellular communication by transferring material between cells.

**Figure 5:**
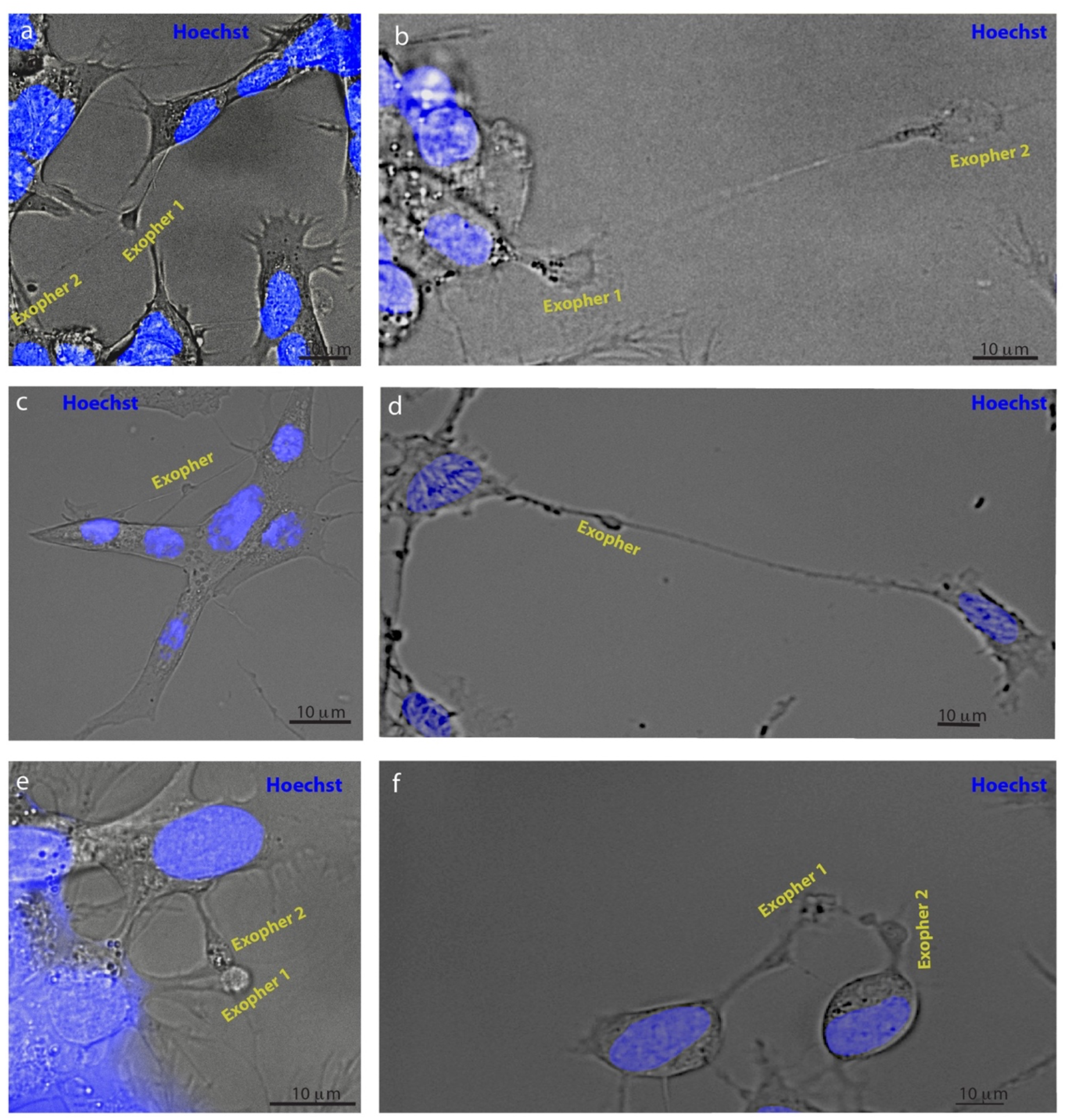
Exophers appear to mediate neuronal communication. SH-SY5Y cells were stained with Hoechst and imaged using brightfield and fluorescence microscopy. a, b) Examples of exophers emanating from another exopher (N = 4 observations). c, d) Examples of exophers connected to two or more different cells via nanotubes (N = 6). e, f) Examples of exophers originating in two adjacent cells in contact with each other (N = 3).

Intriguingly, though in most cases each exopher emanated from a single cell, we also found rare cases in which a single cell had two exophers attached. Although these were uncommon observations, we found such double-exopher cells in all the systems examined, including human brain, (AD2 sample, Supplemental Fig. 6a), wild-type mouse hippocampal neurons (Supplemental Fig. 6b), and SH-SY5Y cells (Supplemental Fig. 6c).

To gain additional insight into exopherogenesis, we performed time-lapse imaging of the SH-SY5Y cells, taking images every 15 min for 12 h. To avoid damaging the cells by repeated laser irradiation and facilitate observation of nanotubes, we used brightfield images again in these experiments. Because exopherogenesis is a rare event and no specific markers currently are available, we could not observe the initial budding of exophers and instead followed exophers identified after their initial formation. The time-lapse images showed that during the experiment, the exopher size changed dynamically. Two exophers were observed independently for 12 h. Images are shown only for exopher 1 (Supplemental Fig. 7a-l and Supplemental Movie 2). Both exophers appeared to increase in size over time and then decrease Supplemental Fig. 7m, but the kinetics and sizes were distinct.

Previously, the presence of cellular organelles, including mitochondria and lysosomes, has been reported in *c. elegans* and mammalian cardiomyocyte exophers^1,9^. To test whether mammalian neuronal exophers also contain cellular organelles, we stained SH-SY5Y cells using markers of mitochondria (MitoTracker™ Deep Red FM, Supplemental Fig. 8a), autophagosomes (Premo™ Autophagy Sensor LC3B-GFP, Supplemental Fig. 8b), endoplasmic reticulum (ER-Tracker™ Green, BODIPY™ FL Glibenclamide, Supplemental Fig. 8c), lysosomes (LysoTracker™ Deep Red, Supplemental Fig. 8d), and Golgi (CellLight™ Golgi-GFP, BacMam 2.0, Supplemental Fig. 8e). We analyzed 20-23 exophers in each case and found each of the organelles in most (78 ± 5%) of the exophers.

Thinking about the mechanism by which cells produce exophers emanating from their plasma membrane, we hypothesized that cytoskeletal proteins must participate in the process. Staining SH-SY5Y cells for cytoskeletal proteins, we found that in addition to β-III tubulin and MAP-2, actin is also prevalent in the nanotubes and exophers, as might be expected. Interestingly, although in many cases the fluorescence intensity of β-III tubulin (Supplemental Fig 9a, b), MAP-2 (Supplemental Fig 9c, d), and actin (Supplemental Fig e, f) in the exophers was similar to that of the parent cell, we also observed certain instances in which the intensity of each of these cytoskeletal proteins staining was substantially lower, or higher, than in the parent cell, suggesting that cells may use different mechanisms to achieve exopherogenesis.

## Discussion

Eukaryotes have developed several processes for managing the turnover of cellular components. The two most common mechanisms, UPS and ALP, have been studied extensively, whereas removal of cellular components via exophers is a newly discovered process^1,7–9^. Here, we show for the first time that this mechanism is conserved in mammalian neurons, including those originating from mice and humans. Similar to the previous findings in nematodes and mammalian cardiomyocytes, we found that exophers form by evagination of the cell membrane and are connected initially to the originating cell via a nanotube, until eventually they appear to disconnect from the cell body. In addition, as was reported previously, we observed multiple types of organelles in exophers^1,9^.

Our findings suggest that in addition to their roles in removal of unwanted cellular components^1,9^ or supporting nematode embryos^8^, a basal, innate level of exopher formation exists in young, wild-type mouse neurons (Fig. 3d) and human neuroblastoma cells (Supplemental Fig. 5), and that these exophers tend to have a relatively small area, ~10 μm^2^. The observations of such exophers connected to two cells (Fig. 5c, d) or extended to a relatively large distance (Fig. 5a, b) suggest that they serve as a mechanism for intercellular transfer of cargo that is too large to pass through a direct nanotube. This innate exopherogenesis is reported here for the first time.

We also found that when neurons are exposed to proteostatic stress, such as elevated concentrations of wild-type or mutant human tau, the number of exophers increases (Fig. 2d, Fig. 3, Supplemental Fig. 1d), suggesting an adaptive response. The adaptive exophers containing pathological tau tend to have a larger area compared to the innate exophers (Fig. 4). These findings support the hypothesis of Driscoll and co-workers based on their findings in *c. elegans*^1,7^ of the role of exophers as an adaptive response to proteostasis and reveal that this response is conserved in mouse and human neurons.

Melentijevic et al. reported high size variability of exophers in the *c.elegens*, from 0.8 to 47.8 μm^2^, and an average size of 11.3 μm^2^. We found a similar size variability in the mammalian neurons we studied, though the median size and size range were similar in all systems: human brain – 1.4 to 253.9 μm^2^, median 27.5 μm^2^ (95% CI, 19.1 to 44.4) primary neurons – 3.3 to 245.4 μm^2^, median 27.2 μm^2^ (95% CI, 12.6 to 47.6) SH-SY5Y – 1.5 to 204.12 μm^2^, median 25.6 μm^2^ (95% CI, 13.2 to 57.5). The median size and size range of mammalian exophers were considerably larger than those reported previously in *c.elegans*, likely due to the smaller size of *c. elegans* and their neurons (5-10 μm^2^ soma size)^17^ compared to mammalian neurons (5-100 μm^2^ soma size). Interestingly, in *c. elegans* the size of muscle-cell exophers, 2–15 μm in diameter, was in the same range is the average neuronal-exopher diameter, whereas the reported average diameter of mammalian cardiomyocyte exophers, 3.5 ± 0.1 μM^9^, is substantially smaller and more homogeneous than the median diameter and wide range of diameters of mammalian neuronal exophers.

In the absence of unique markers, our identification of exophers was based on their morphology and the absence of nuclear staining. The reliance on morphology may raise the concern of mixing exophers with similar structures, such as blebbing. Blebbing is a common phenomenon involved in cytokinesis, cell spreading, and locomotion, yet it has been associated mostly with cell injury and apoptosis^18^. Although morphologically blebbing also constitutes protrusions of the plasma membrane toward the extracellular space, similar to exophers, these protrusions typically have an area of ~2 μm^2^ and are short-lived, expanding for ~30 seconds and retracting for ~120 seconds^19^, distinguishing them from exophers.

Our study opens many questions about exopher neurobiology. What specific mechanisms control their formation? Are they similar to those reported in nematodes? What cell-autonomous and non-cell-autonomous signals lead to exopherogenesis in the brain? Which genes are involved? Are there signals on the surface of exophers directing them to other neurons or to glial cells as content carriers? Is there an “eat me” signal for adaptive neuronal exophers, akin to those reported for cardiomyocytes signaling to macrophages^9^, after they disconnect from the originating cells? Do neuronal exophers also signal to and interact with astrocytes? We hope that the discoveries reported here will prompt future exploration of the basic biology of exophers and their roles both in normal physiology and under disease and injury conditions.

## Methods

### Cryosectioning of human brains

Frozen hippocampi from two healthy controls, two patients with AD, one patient with FTLD, and one patient with PSP (Table 1) were sectioned using a Leica CM3050 S cryostat into 30-μm sections, which were fixed immediately in 4% (w/v) paraformaldehyde for 30 min at room temperature. The sections were stored at 4 °C in 0.1M phosphate-buffered saline (PBS) containing 0.05% sodium azide until use.

### Mouse models

P301S-tau (Tg (Prnp-MAPT*P301S)PS19Vle/J), hTau (B6.Cg-*Mapt*^tm1(EGFP)Klt^ Tg(MAPT)8cPdav/J) transgenic mice, and C57BL/6J wild-type mice were obtained from Jackson Laboratories and bred and maintained by the UCLA Division of Laboratory Animal Medicine. Homozygous P301S-tau, hTau, and wild-type littermates were used in this study. All the experiments were reviewed and carried out following National Research Council Guide for the Care and Use of Laboratory Animals approved by the University of California-Los Angeles Institutional Animal Care Use Committee and performed with strict adherence to the guidelines set out in the National Institutes of Health Guide for the Care and Use of Laboratory Animals. Mice were anesthetized using isoflurane and perfused transcardially using 0.1 M PBS, pH 7.4, containing 1% protease and phosphatase inhibitors (Sigma-Aldrich). Brains were removed and fixed immediately by immersion in 4% (w/v) paraformaldehyde in 0.1 M PBS for immunohistochemistry.

### Immunohistochemistry of mouse brain sections

Free-floating sections were immersed in antigen-retrieval buffer comprising 10 mM sodium citrate, 0.05% (v/v) Tween-20, pH 6.0, and incubated for 30 min at 60 °C. Then, the sections were immersed in PBS containing 0.1% (v/v) Triton-X100 (PBS-T) for 15 min at room temperature for permeabilization. The sections were blocked using 5% (w/v) bovine serum albumin (BSA) in PBS at room temperature for 60 min and incubated with primary antibodies (Table 3) in 2.5% (w/v) BSA in PBS overnight at 4 °C with gentle agitation. The sections were washed thrice in PBS containing 0.03% (v/v) Tween-20 (PBS-T) for 10 min and subsequently incubated for 2 h at room temperature in the dark with secondary antibodies diluted in 2.5% (w/v) BSA in PBS (Table 3). The sections again were washed thrice and then mounted onto gelatin-coated slides using ProLong™ Diamond Antifade mountant with DAPI (Invitrogen-ThermoFisher). Fluorophore-conjugated antibodies were visualized using a Leica confocal SP8 microscope.

**Table 3.** Antibodies used in the project.

| **Table 3. Antibodies used in the project** |  |  |  |  |  |  |
| --- | --- | --- | --- | --- | --- | --- |
| **Target** | **Primary Antibody** | **Dilution** | **Source** | **Secondary antibody** | **Dilution** | **Source** |
| p-Tau | AT8 | 1:200 | Thermo Fisher | AlexaFluor555 (anti-mouse) | 1:200 | Thermo Fisher |
| Microglia | Anti-Iba1 | 1:100 | Abcam | AlexaFluor647 (anti-rabbit) | 1:200 | Thermo Fisher |
| Total-Tau | HT7 | 1:200 | Cell Signaling | AlexaFluor595 (anti-mouse) | 1:200 | Thermo Fisher |
| Aβ42 | H31L21 | 1:100 | Invitogen | AlexaFluor647 (anti-rabbit) | 1:200 | Thermo Fisher |
| MAP-2 | Anti-MAP-2 | 1:200 | Synaptic System | AlexaFluor594 (anti-guinea pig) | 1:200 | Synaptic System |
| β-III tubulin | Anti-β-III Tubulin | 1:200 | Synaptic System | AlexaFluor488 (anti-chicken) | 1:200 | Thermo Fisher |

### Immunohistochemistry of human brain sections

Free-floating sections were stained exactly following the same protocol used for the mouse brain section staining described above. Fluorophore-conjugated antibodies were visualized using a Keyence BZ-X710 fluorescence microscope.

### Cell Culture

Primary mouse hippocampal neurons were derived from postnatal day 1 wild-type, P301S-tau, or hTau mice. Isolated hippocampi were incubated in a 0.5-mg/mL papain and 0.6 μg/mL DNAase solution for 20 min at 37 °C. Then, the tissue was washed and triturated by pipetting 20 times through filtered 1000-μL tips. Dissociated cells were collected by centrifugation (200 *g*, 3 min at 25 °C) and resuspended in complete medium comprising neurobasal medium (Gibco, 21103) supplemented with 2% B-27 plus (Gibco A3582801), Antibiotic-Antimycotic (100 units/mL of penicillin, 100 μg/mL of streptomycin and 250 ng/mL of amphotericin B, Gibco 15240062), and GlutaMax. Cells from both hippocampi of each mouse were pooled together and plated in 2 wells of an 8-well Chambered Coverglass (ThermoFisher, 155409PK) at a density of ~250,000 cells per chamber. The coverslips were coated with 0.5-mg/mL poly-ornithine (Sigma, P8638), followed by coating with 5 μg/mL laminin (Corning, 354232) at room temperature (RT) for 16 h. Non-neuronal cells were removed by addition of 2 μM cytosine arabinoside (Acros Organics, AC449560010) after 72 h in culture. Antibiotics-depleted complete medium was replenished twice weekly, replacing 1/3 of the volume each time. Cells were inspected manually at 60X magnification for the presence of exophers on day 7, 14 and 21 of the culture.

Undifferentiated human neuroblastoma SH-SY5Y cells were cultured in DMEM/F12 medium (Gibco, 11320033) supplemented with 10% fetal bovine serum (FBS) and a penicillin-streptomycin solution (100 units/mL of penicillin and 100 μg/mL of streptomycin, Caisson labs, PSL01). SH-SY5Y cells were plated at 30,000 cells per well in 8-chamber slides and grown to 60% confluency.

### Immunocytochemistry

Primary neurons were fixed in 4% (v/v) paraformaldehyde in PBS for 15 min, washed in PBS thrice, and permeated for 1 h at room temperature in blocking buffer containing 0.1% BSA, 5% donkey serum, 0.3% Tween-20 in PBS, pH 7.4. The blocking buffer then was removed, and the cells were stained with anti-human tau monoclonal antibody HT7 (Invitrogen) at 1:1,000 dilution in blocking buffer overnight at 4 °C, followed by Alexa-Fluor-555-conjugated donkey anti-mouse secondary antibody (Thermo Fisher Scientific) at 1:500 dilution in blocking buffer for 1 h at room temperature. After washing in PBS, the coverslips were mounted on microscope slides using Prolong Gold Antifade reagent with DAPI (Thermo Fisher Scientific) and imaged using a Leica SP8-SMD Confocal Laser Scanning Microscopy Platform.

SH-SY5Y cells were grown to 60% confluency and incubated with markers of mitochondria (MitoTracker™ Deep Red, Invitrogen), lysosomes (LysoTracker™ Deep Red, Invitrogen), autophagosomes (Premo™ Autophagy Sensor LC3B-GFP, ThermoFisher), endoplasmatic reticulum (ER-Tracker™ Green (BODIPY™ FL Glibenclamide), Fisher Scientific), or the Golgi system (CellLight™ Golgi-GFP, Thermo Fisher). After 16 h of incubation, Hoechst dye (Invitrogen) was added for 30 min. Images were taken at 60X magnification. 3D movies were generated from Z-stack images using the real-time 3-D image viewer of BZ-X Analyzer software.

### Live cell Imaging

SH-SY5Y cells were cultured in Chambered Coverslips at ≤ 60% confluence. Hoechst dye was added 30 min before the beginning of the imaging. Cells were brought into focus at 60X magnifications and exophers were sought manually. Exophers were brought into focus and live-cell, time-lapse images were taken using a BZ-X710 fluorescence microscope (Keyence) at 15-min intervals for 12 h. Movies were created by combining all images using the video making option of BZ-X analyzer software. Images were analyzed using BZ-X Analyzer (Keyence) and ImageJ.

## Statistical analysis

We used one-way or two-way analysis of variance (ANOVA), as appropriate, for analysis of cell culture data in Prism 9.0 (GraphPad, La Jolla, CA). The Poisson regression model was used to evaluate the expected number of exophers per cm^2^ in the disease human brains compared to the control brains. To account for differences in the areas of different sections, we included area as an exposure variable to this model. Multiple testing adjustments were performed using the FDR criterion. P-values were adjusted for multiple comparisons using Hochberg criteria to control the family-wise error rate at < 5% (3 pre-specified comparisons).

## Acknowledgments

We thank Dr. David Teplow for helpful comments on the manuscript. The work was supported by NIH/NIA grants R01AG050721 and RF1AG054000.

