## Supplemental Information for "Exophers are components of mammalian cell neurobiology in health and disease"

**Supplemental Figures and Legends**

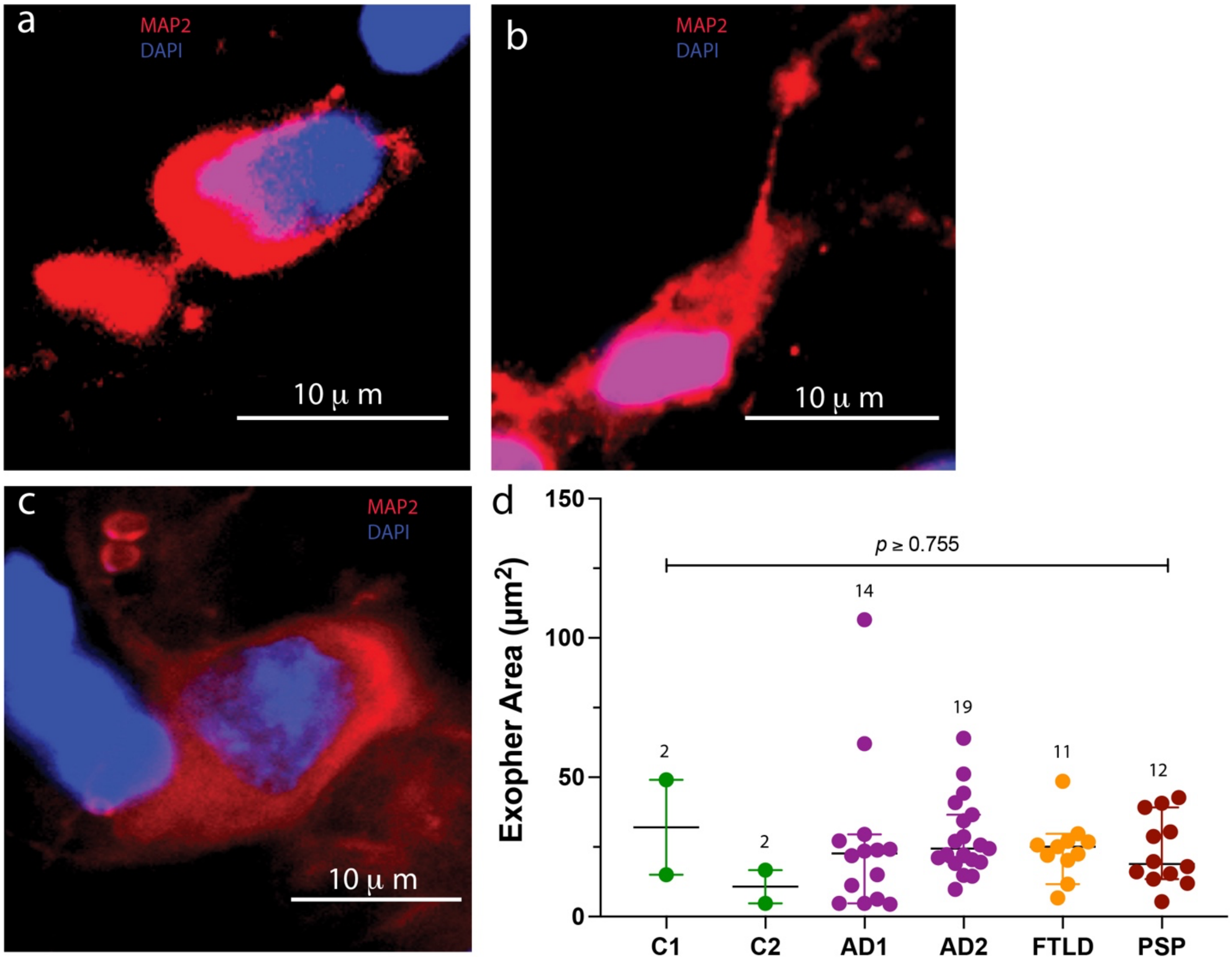


**Supplemental Fig. 1. Human-brain exophers.** Human hippocampal sections were stained with an anti-MAP-2 antibody (red) to outline the morphology of the cells and exophers and nuclei were

stained with DAPI (blue). a) An exopher connected directly to the cell body. b) An exopher connected via a nanotube. c) Exophers separated from the adjacent cell. d) Comparison of exopher areas in human brains. The data are presented as median ± CI . The number of exophers observed in each case is indicated above each column. *P*-values were calculated using a one-way ANOVA with post-hoc Tukey test.

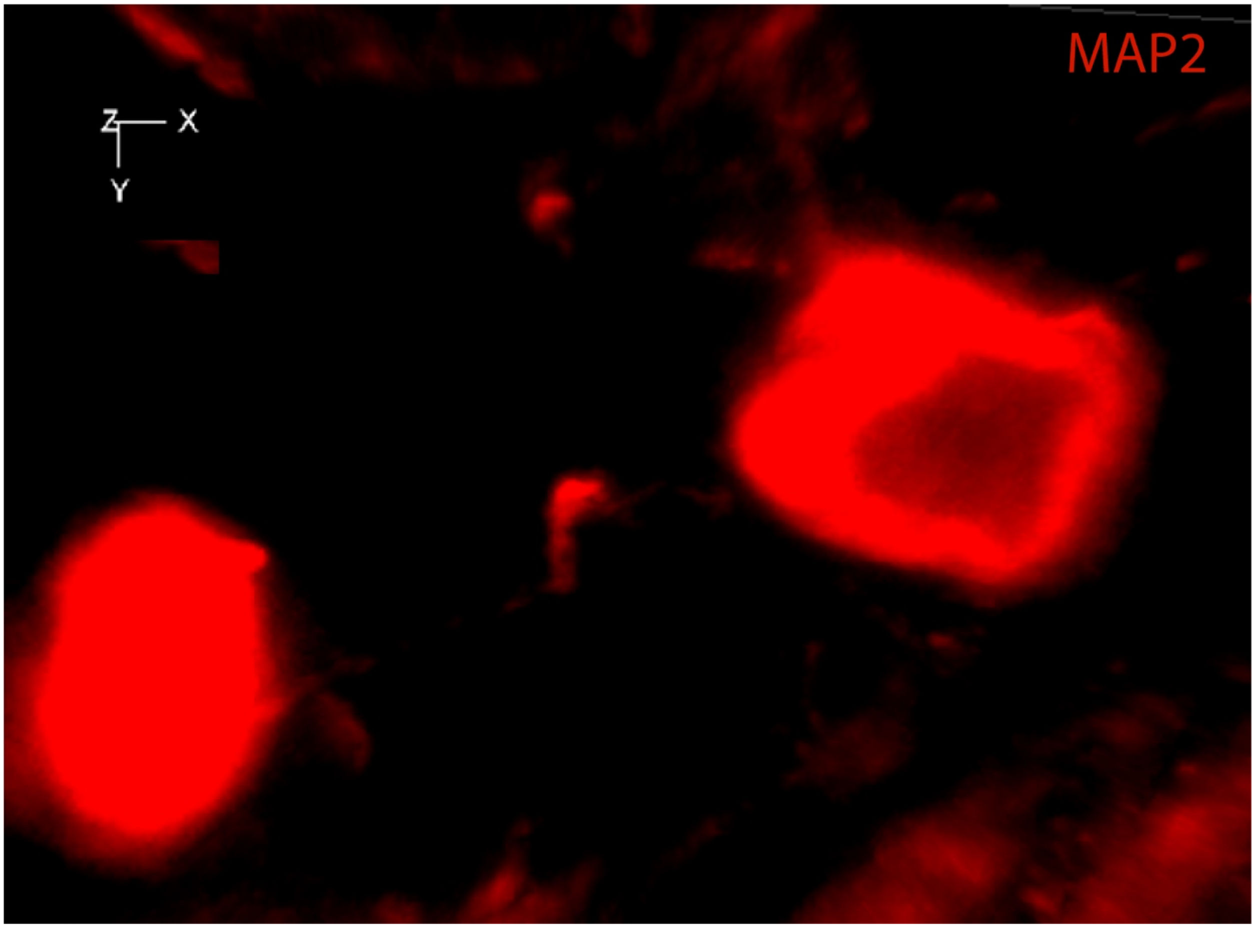


**Supplemental Fig. 2: A small exopher midway between a cell and a large exopher.** A FTLD human brain was stained for MAP-2 and DAPI. This image is one frame from Supplementary Movie 1. The software does not allow showing two channels in movies and therefore the DAPI staining is not shown and the nucleus appears as a dark shadow.

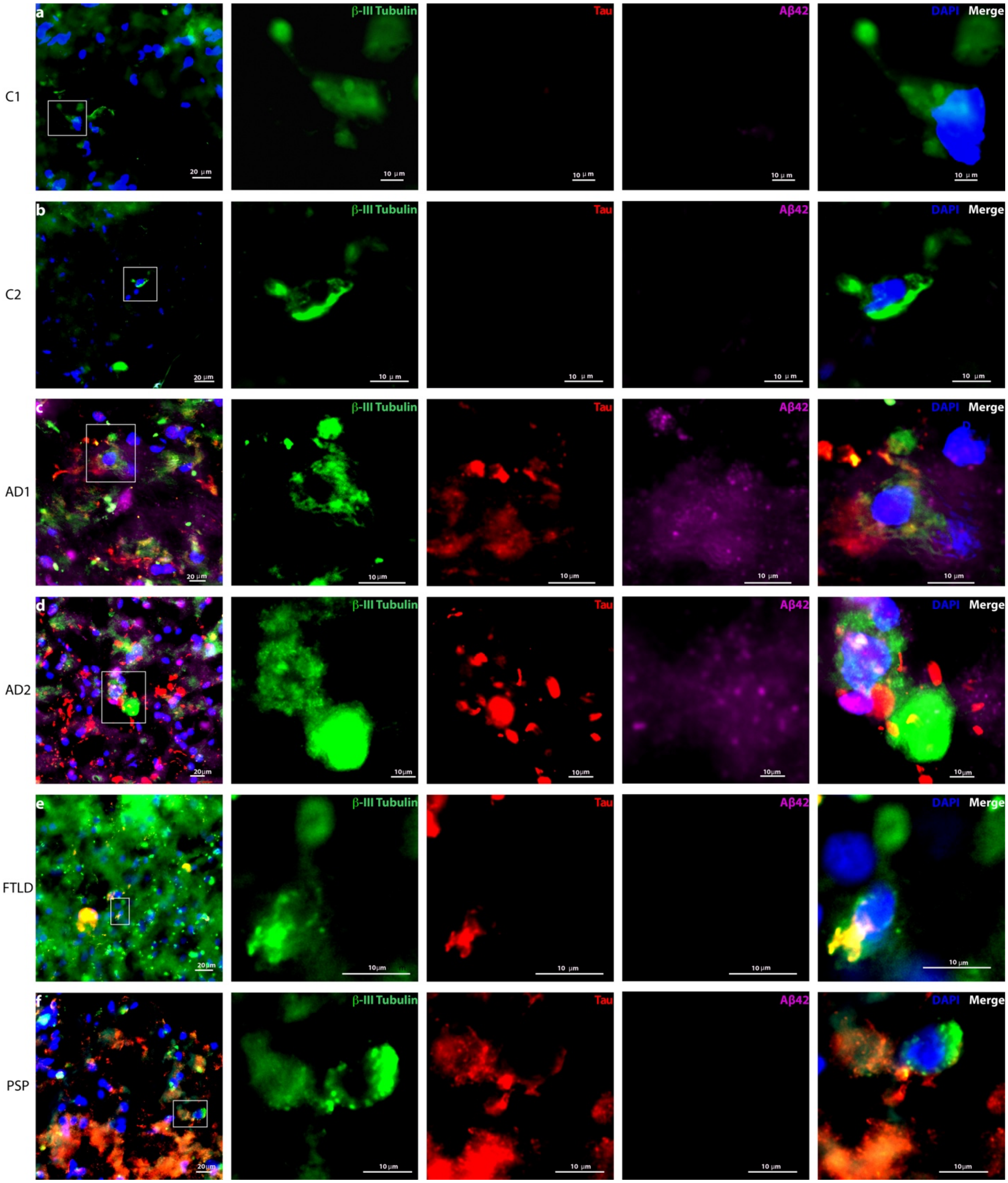


**Supplemental Fig. 3. Analysis of Aβ and p-tau in human-brain exophers.** Brain sections from each human brain were stained for β-III tubulin (green), p-tau (AT8, red) and Aβ42 (H31L21, magenta) and DAPI (blue). First image of each row is a broad view of the section, in which the zoomed-in area in the subsequent images is highlighted by a white rectangle.

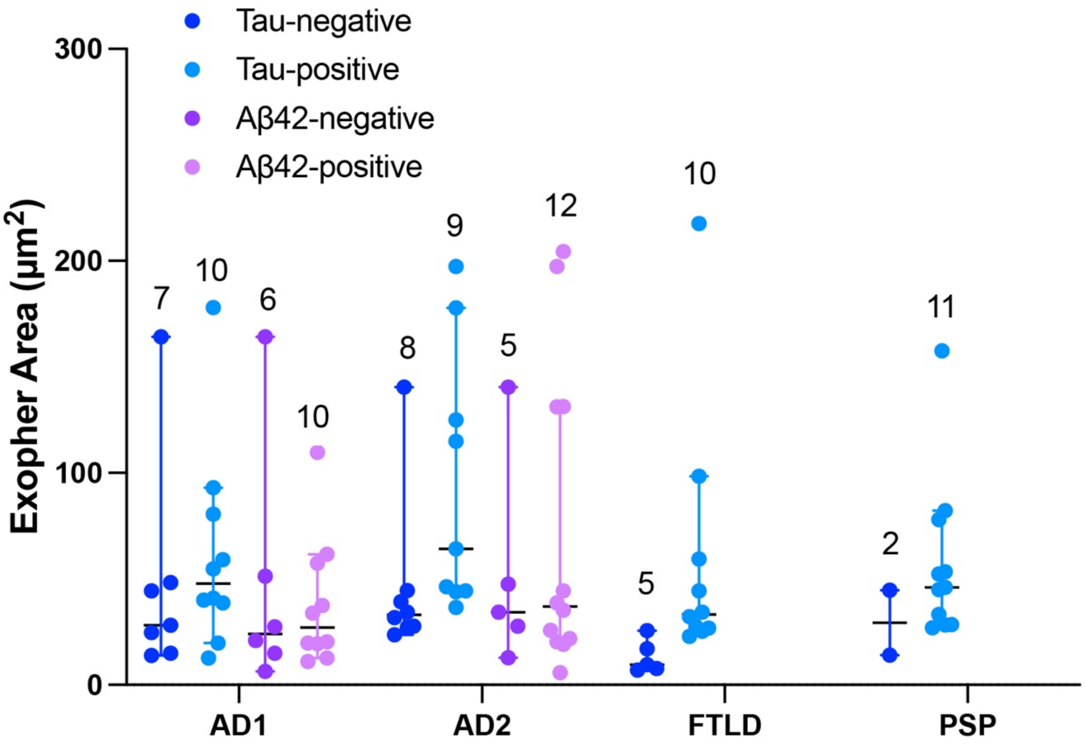


**Supplemental Fig. 4. Quantitative analysis of human brain exophers stained for Aβ42 and p-tau.** Exopher areas in human brain sections stained with antibodies against Aβ42 and p-tau were measured using ImageJ. The number of exophers observed in each case is indicated above each column. *P*-values were calculated using a two-way ANOVA with post-hoc Tukey test.

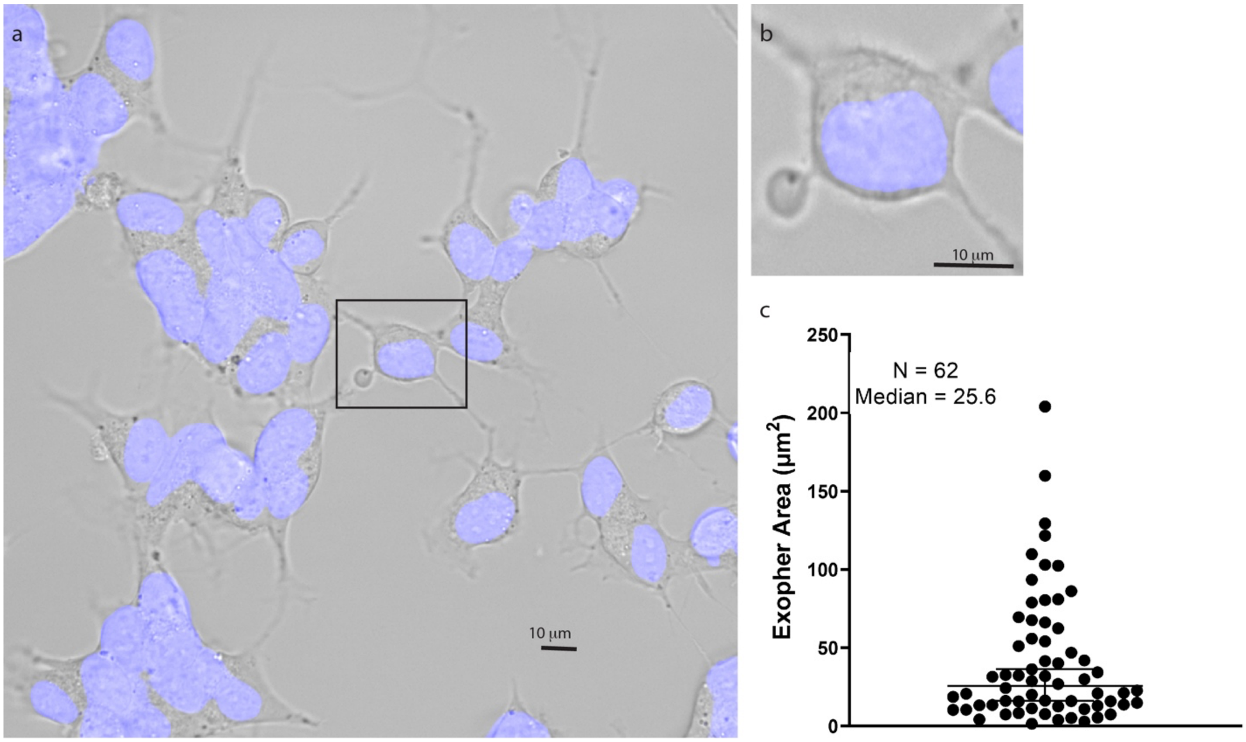


**Supplemental Fig.5. Exophers in SH-SY5Y cells.** a) A representative image of the SH-SY5Y cell culture. A cell with an emanating exopher is highlighted in the center of the image and this area is enlarged in panel b for better visualization of the nanotube. c) The area of exophers found in the SH-SY5Y cells was quantified using ImageJ.

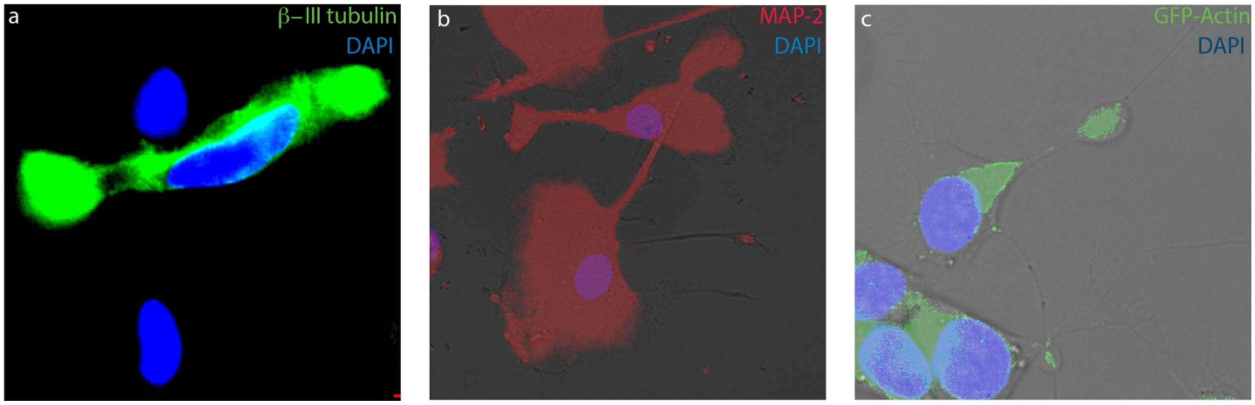


**Supplemental Fig.6. Examples of two exophers attached to the same cell.** Cells attached to two exophers were found rarely in the different systems analyzed. a) A cell attached to two exophers in a human brain stained with βIII-tubulin and DAPI (N = 2 out of in 158 exophers analyzed). b) A cell attached to two exophers in primary hippocampal wild-type mouse culture stained with MAP-2 and DAPI ((N = 2 out of in 499 exophers analyzed including both P301S-tau and hTau mice). c) A cell attached to two exophers in SH-SY5Y cells stained with Hoechst and expressing with GFP-Actin (N = 2 out of in 125 exophers).

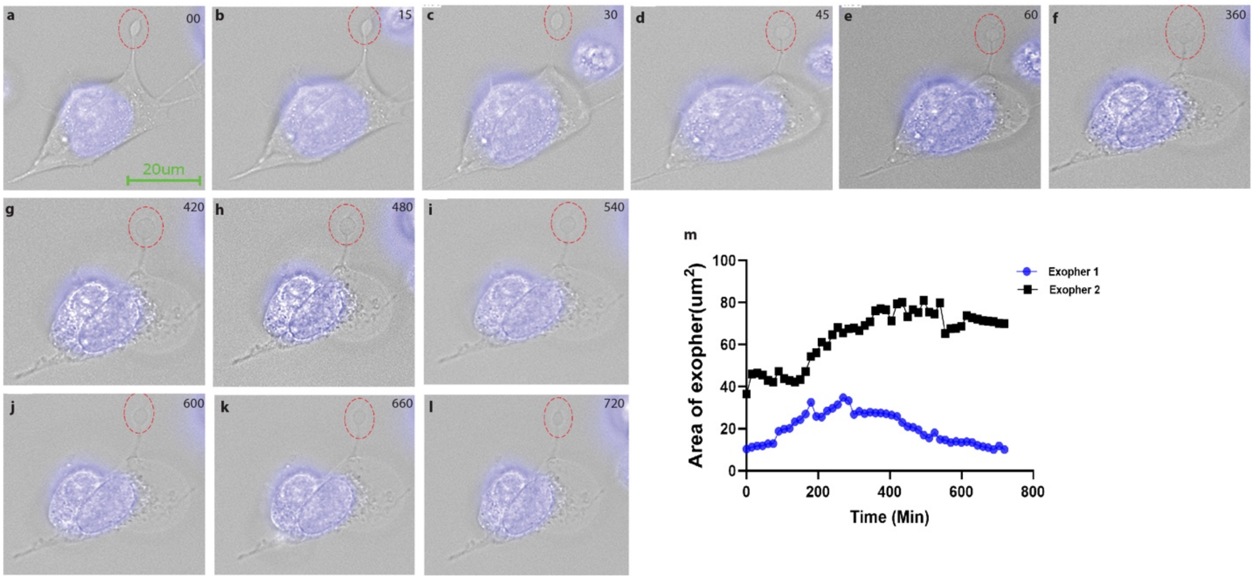


**Supplemental Fig.7. Time-lapse imaging of exophers.** a-l) Images of a SH-SY5Y cell bearing an exopher between 0–720 min. m) Temporal change in the area of two independent exophers during 720 min of time-lapse imaging.

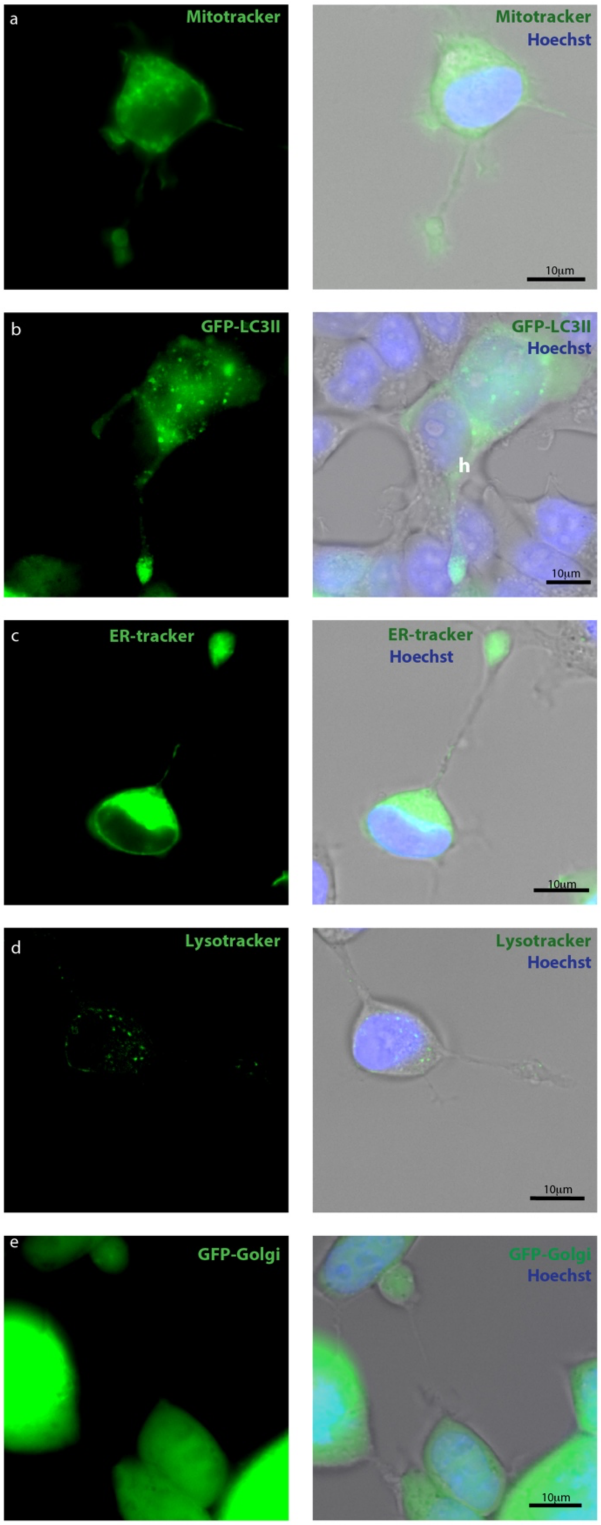


**Supplemental Fig. 8. SH-SY5Y-cell exophers contain various cellular organelles:** SH-SY5Y cells were stained with markers for different organelles (pseudo-colored green in all cases) and

Hoechst (blue) for visualization of nuclei. The fluorescence image of the organellar marker is shown on the left and the overlap with Hoechst and the brightfield image on the right. a) MitoTracker™ Deep Red FM for staining of mitochondria, 17 out of total 20 exophers were stained positive. b) Premo™ Autophagy Sensor LC3B-GFP for staining of autophagosome, 15 out of 20 exophers were stained positive for autophagosomes. c) ER-Tracker™ Green (BODIPY™ FL Glibenclamide for staining of endoplasmic reticulum, 17 out of 21 exophers were found positive for ER. d) LysoTracker™ Deep Red for staining of lysosomes, 16 out of 22 exophers were found positive for lysosomes. e) CellLight™ Golgi-GFP for staining of the Golgi apparatus, 18 out of 23 exophers were found positive for Golgi staining.

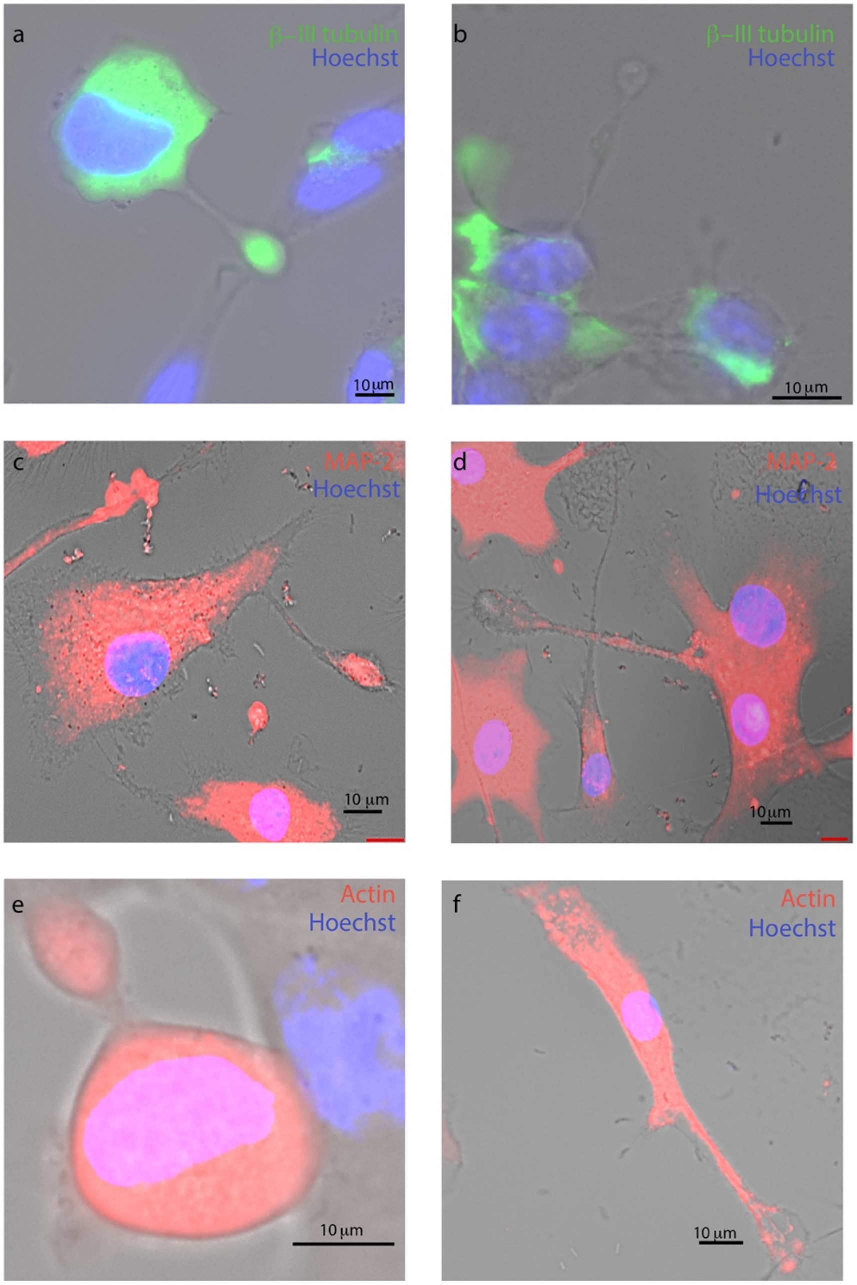


**Supplemental Fig. 9: Exophers contain inconsistent amounts of β-III tubulin, MAP-2, and actin.** SH-SY5Y cells were fixed and stained for β-III tubulin (a, b) or MAP-2 (c, d). Other cells expressed transiently red fluorescent protein-conjugated actin (CellLight$^{TM}$Actin-RFP). In each pair, the images were chosen to demonstrate the inconsistent level of the respective structural protein in the exopher compared to the parent cell.
